# Ultraconformable cuff implants for long-term bidirectional interfacing of peripheral nerves at sub-nerve resolutions

**DOI:** 10.1101/2023.04.14.536862

**Authors:** Alejandro Carnicer-Lombarte, Alexander J. Boys, Amparo Güemes, Johannes Gurke, Santiago Velasco-Bosom, Sam Hilton, Damiano G. Barone, George G. Malliaras

## Abstract

Implantable devices interfacing with peripheral nerves exhibit limited longevity and resolution. Poor nerve-electrode interface quality, invasive surgical placement and development of foreign body reaction combine to limit research and clinical application of these devices. Here, we develop cuff implants with an ultraconformable design that achieve high-quality and stable interfacing with nerves in chronic implantation scenarios. When implanted in sensorimotor nerves of the arm in awake rats for 21 days, the devices recorded nerve action potentials with fascicle-specific resolution and extracted from these the conduction velocity and direction of propagation. The ultraconformable cuffs exhibited high biocompatibility, producing lower levels of fibrotic scarring than clinically equivalent PDMS silicone cuffs. In addition to recording nerve activity, the devices were able to modulate nerve activity at sub-nerve resolution to produce a wide range of paw movements. The developed implantable devices represent a platform enabling new forms of fine nerve signal sensing and modulation, with applications in physiology research and closed-loop therapeutics.

---

The peripheral nervous system (PNS) is formed by a vast signal-transmitting network linking the central nervous system (CNS) with most organs and structures in the body. Nerves provide the CNS with information on the state of the structure to which they connect and transmit modulation signals back to the periphery. This wide anatomical distribution and unique physiological role makes nerves an attractive target for interfacing with implantable devices^1–3^. Nerve implants can be deployed to a particular nerve to gain information on the physiological state of the organ it innervates via electrical recording, or to modulate function of that organ through electrical stimulation. This flexibility has allowed their use in a multitude of applications^4–7^.

While the research and treatment opportunities offered by nerve interfacing are substantial, translation into applications has remained limited due to the inadequate long-term performance of most interfaces^8^. Implantable nerve interfaces are generally classified based on their location in or around nerves. Low-invasive epineurial interfaces – usually in the form of cuffs – wrap around the circumference of the nerve and interface using one or few electrodes providing low selectivity recording and stimulation capabilities^1,8^. Cuffs are used in the clinic for whole-nerve stimulation therapies such as vagus nerve stimulation^6,7^. However, most nerves are composed of fascicles connecting to multiple targets, making whole-nerve interfacing unsuitable for many recording and stimulation applications. As an alternative to cuffs, intraneural or penetrating nerve implants pierce the nerve epineurium and deploy electrodes throughout the endoneurium^1^. Their more invasive design allows for the recording and stimulation of select portions of the nerve, enabling a wider range of applications. However, the higher tissue damage caused by penetrating devices and their reliance on a close interface with axons makes them highly vulnerable to chronic inflammation and foreign body reaction (FBR) – a slow-developing process affecting both cuffs and penetrating implants leading to their envelopment in a fibrotic scar^8^. FBR and tissue-implant interface degradation limit long-term use of penetrating nerve interfaces, particularly for recording applications^2,8^.

Advances in materials and microfabrication strategies have opened new opportunities to interface with the nervous system. The last decade has seen the development of a wide range of multielectrode array implants interfacing with nervous tissue in novel ways^9^, with a growing focus on soft and flexible constructions to improve tissue interface quality and long-term stability^9–14^, as well as high-performance recording microelectrode materials^11^. However, most of these technologies have focused on the CNS. Peripheral nerve interface designs have benefitted comparatively less from advances in fabrication. While penetrating nerve interfaces with flexible materials and microelectrode arrays have been developed^15,16^, nerve cuffs in particular have seen little innovation, with many applications continuing to rely on few large electrodes and rigid thick-film-fabricated materials^7^. This has limited their use in long-term applications despite their more clinically-translatable low-invasive design.

In this work we present an implantable ultraconformable nerve cuff which exhibits long-term tissue compatibility and stable recording of nerve activity for longer periods of time. We show these cuffs to achieve sub-nerve recording and stimulating resolutions typically limited to more invasive penetrating implants, as well as the ability to identify axon conduction velocity and direction in awake animals.

## Results

### Design and fabrication of ultraconformable cuffs

We fabricated our implantable cuffs using thin (4 μm) parylene C (PaC) as a substrate. This rendered the devices highly flexible, allowing them to conform well to curved surfaces without damaging them (Fig. 1a). Wiring connections to the poly(3,4-ethylenedioxythiophene):poly(styrenesulfonate) (PEDOT:PSS) microelectrode array were carried by a narrow PaC connector. The shape of this connector was designed to accommodate the anatomy of the rat forelimb, and included suture loops and surgical guidance tabs to facilitate handling and secure the cuffs after implantation (Fig. 1b). Connections were then carried to a flat flexible cable (FFC) connector.

**Fig. 1.**
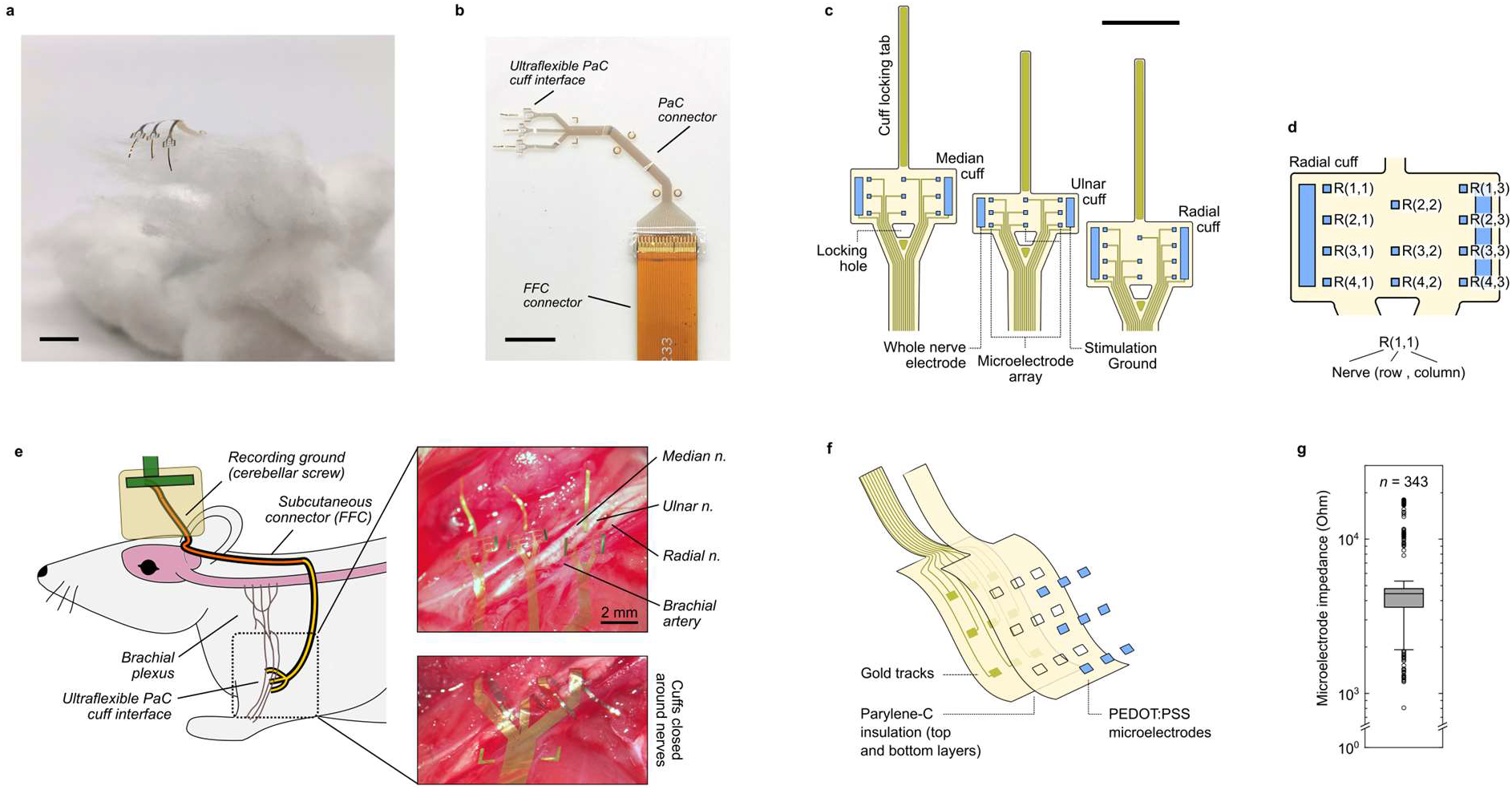
Overview of ultraconformable nerve cuff arrays. **a**, Photograph ultraconformable nerve cuff array device on a cotton ball, highlighting its flexible and lightweight construction. **b**, Photograph of ultraconformable nerve cuff array device. The cuffs are fabricated from parylene C (PaC). These carry electrode connections through a PaC connector, bonded to a flat flexible cable (FFC) connector through which they are externalised. The shape of the PaC connector is designed to accommodate the shape and movement of the rat shoulder, and contains multiple tabs to facilitate surgical manipulation and suturing. **c**, Diagram of cuff and multielectrode array layout. Each implantable device consists of three cuffs designed for implantation into three individual nerves (median, ulnar, and radial nerves). Each cuff consist of a regular array of 100 by 100 µm microelectrodes which are wrapped around the circumference of the nerve during implantation. To seal each cuff, its locking tab is fed through the cuff’s locking hole. In addition to the microelectrodes, each cuff contains a larger electrode which can be used for whole nerve stimulation/recording and a ground used for nerve stimulation. **d**, Diagram displaying naming convention of microelectrodes (radial nerve cuff shown). Note the radial nerve cuff lacks a R(1,2) microelectrode due to device channel number limitations. Microelectrodes within the same column in the array are deployed as a ring around the cuffed nerve when implanted, and are referred to as rings instead of columns in *in vivo* experiments. **e**, Diagram of chronic implantation model and images of device implantation into nerves of the rat forearm. Implanted cuffs microelectrode connections are externalised via a subcutaneously-positioned FFC, exiting the rat of the body through a headcap port. For recording, a ground is implanted into the CSF above the rat cerebellum. **f**, Construction of ultraconformable nerve cuff array device. The device consists of gold tracks encapsulated between two layers totalling to 4 µm of PaC. The tracks connect to PEDOT:PSS microelectrodes. **g**, Impedance of device microelectrodes following fabrication (*n* = 343 microelectrodes, *N* = 12 devices). Scale bar a-b: 10 mm, c: 2 mm.

Our implants were designed with a triple cuff construction to interface with three key nerves of the brachial plexus: the median, ulnar and radial nerves (Fig. 1c). These three nerves are responsible for movement and sensation in the hand of both humans and rats^17,18^, and are being investigated as interfacing targets to restore hand function^15,19^. The three cuffs had a small profile with a width of 2.6 mm each, and lengths of 1.35, 1.0 and 1.6 mm tailored to the circumferences of the three nerves in rats. The cuffs were designed with long gold-reinforced tabs to facilitate implantation and cuff closure through a locking hole at the cuff base (Fig. 1c). Closed cuffs could be sealed by applying a drop of fast-curing silicone at the tip of these tabs, thereby maintaining the cuffs devoid of any sealing materials to provide enhanced conformability to the nerve after implantation. Each cuff contained an array of 100 by 100 μm PEDOT:PSS microelectrodes and two larger electrodes which enveloped the whole nerve – one used to record whole-nerve activity and one used as a ground to allow for bipolar stimulation setups (Fig. 1c-d).

The fabricated designs could be easily implanted into the nerves at the upper portion of forelimbs of rats (Fig. 1e). The high flexibility and low profile of the device allowed cuffs to be implanted into the adjacent median, ulnar and radial nerves without the need for extensive surgical dissection, and could be conformed around shoulder muscles to facilitate normal limb movement post-implantation. Once the device reached the back of the animal, connections on PaC were replaced by the more robust FFC which was capable of supporting the larger displacements present in this part of the anatomy. This allowed the connections to be externalised through a headcap connection port.

The combination of a PaC substrate with gold connections and PEDOT:PSS microelectrodes has been shown to be robust for chronic recording in the more protected environment of the brain^20^. We adapted this microfabricated layer architecture (Fig. 1f) to accommodate the more challenging movement-rich environment of the peripheral anatomy and the forelimb in particular. The low impedance of PEDOT:PSS microelectrodes seen in other applications was also seen in our devices, with median impedance values of 4.43 kOhm (Fig. 1g).

### Long-term tissue compatibility of cuff implants

To determine the suitability of the developed ultraconformable cuffs for long-term applications, we examined the tissue response to them over 28 days of implantation into the radial nerve of rats. We compared the degree of FBR between the nerve epinerium and the inner surface of the PaC cuffs, characterised by αSMA-positive myofibroblasts^21^, to that of 28 day-implanted medical-grade polydimethylsiloxane (PDMS) and polyethylene (PE) cuffs (Fig. 2a). Both of these materials are biocompatible, with PDMS considered the gold-standard for nerve cuff implants due to its low stiffness^7,8,10^. Nerves implanted with both PDMS and PE cuffs developed a ∼25 μm thick fibrotic capsule, with PDMS capsules exhibiting high variability but overall significantly lower αSMA stain than PE ones (Fig. 2b-c). In contrast, PaC cuff-implanted nerves exhibited a consistently near-absent degree of fibrosis with capsule αSMA barely above background stain values (mean capsule-to-background ratio for PaC of 1.08. PDMS: 2.78-fold increase, PE: 4.72-fold increase) and significantly lower than the other cuffs tested (Fig. 2b-c).

**Fig. 2.**
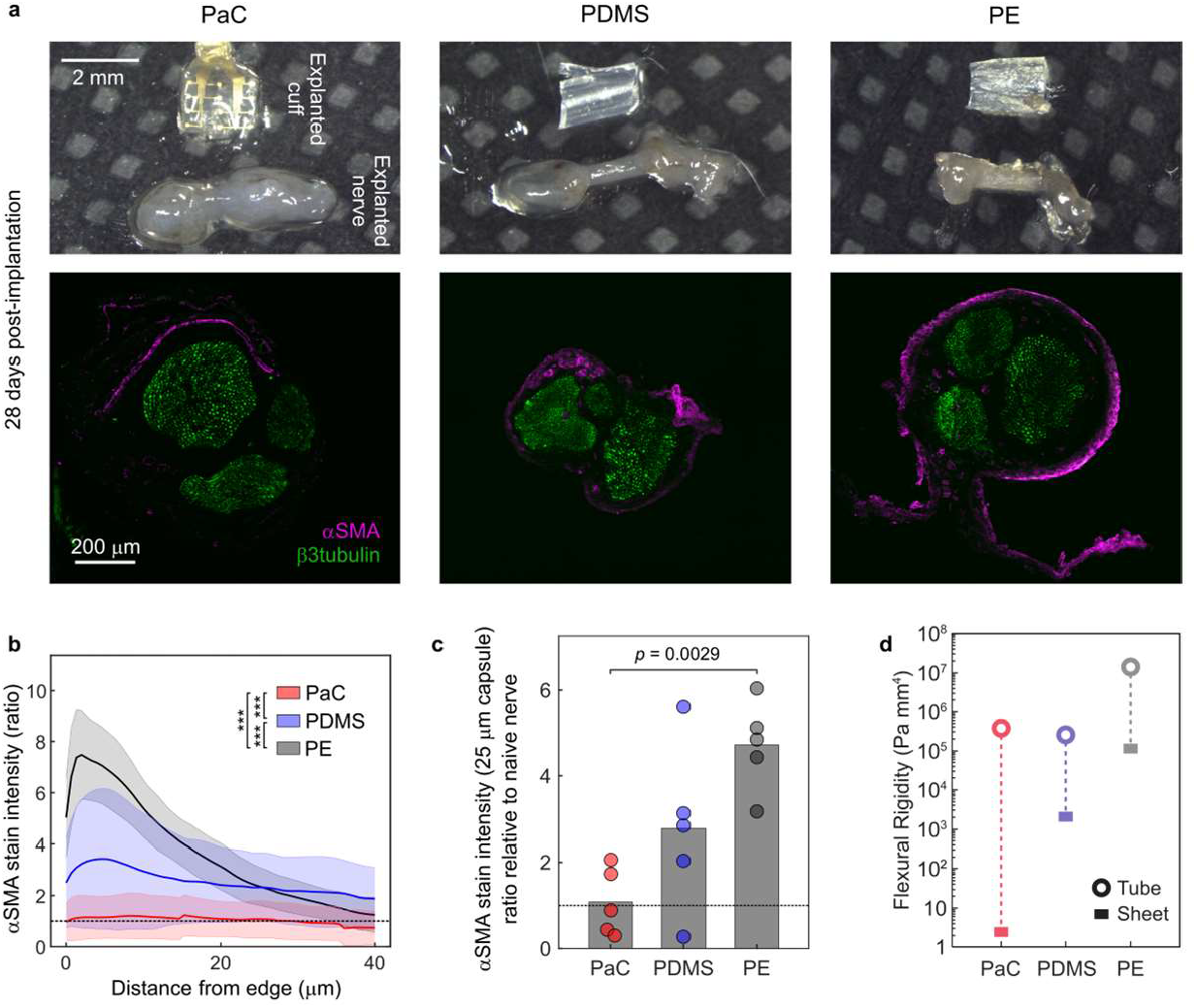
Ultraconformable PaC nerve cuffs exhibit minimal foreign body reaction following four weeks of implantation. **a**, Photographs and immunohistological characterisation of nerve cross-section images of explanted cuffs and radial nerves four weeks after implantation. Nerves implanted with ultraconformable PaC cuffs exhibit little fibrosis (αSMA) when compared to stiff polyethylene (PE) cuffs but also soft silicone (PDMS) cuffs. Axons stained by β3tubulin to show their position relative to the edge of the nerve. **b**, Quantification of fibrosis (αSMA stain) at varying depths along the nerve circumference. Distance from edge represents 0 as tissue in direct contact with the cuff wall. Stain intensity presented as a ratio of stain relative to non-fibrosed tissue at the centre of the nerve. Ultraconformable PaC cuffs cause lower fibrosis than both PE and PDMS cuffs and almost no fibrosis above background levels (deep nerve endoneurium). Lines represent group mean and shaded areas standard deviation. *N* = 5 rats. ***: *p* < 0.001, two-way ANOVA and Tukey post-hoc test. **c**, Quantification of fibrosis (αSMA stain) at the 25 µm of nerve tissue closest to the cuffs. Ultraconformable PaC cuffs cause a significantly lower degree of fibrosis than PE cuffs (*p* = 0.0029, ANOVA and Tukey post-hoc test). *p* values for comparisons with *p* > 0.05 not included in graph. Bars represent the group mean and circles represent values for individual animals. **d**, Flexural rigidity for the different cuffs for a tube (open circle) and for a sheet (rectangle). Dashed line represents the expected range of flexural rigidity between these extremes. Flexural rigidity is calculated as the product of Young’s modulus and moment of inertia for the different shapes. The calculations are detailed in Supplementary Fig. 1. PaC exhibits a lower flexural rigidity range than both PDMS and PE due to its low thickness, providing the cuffs with high flexibility, conformability, and long-term tissue compatibility.

To understand the observed variation in FBR, we examined the mechanics for each cuff type. Mechanical mismatch between implant and tissue is generally recognised as the major driver of FBR and tissue damage, particularly in the nervous system^8,10,22–24^. First, we determined the stiffness for each material (Young’s modulus PaC: 1.13 GPa, PDMS: 2.86 MPa, PE: 217 MPa, n = 4 - 7 samples) (Supplementary Fig. 1a-d), finding no trend with observed FBR. We used these values to calculate the flexural rigidity for each cuff, which also takes geometrical effects into account^25^. Given the wide range of loading conditions that could be present in the complex loading environment for a nerve cuff, we calculated a range of possible flexural rigidities that represent the different extremes (Supplementary Fig. 1e). On one extreme, we assumed the bending of the entire nerve cuff as a tube and on the other extreme, we assumed a rectangular cross-section, representing a point force on a singular small section of the cuff (Fig. 2d). As the nerve has an average stiffness of ∼60 Pa^26^, which results in a flexural rigidity of 0.38 Pa mm^4^ (calculating the moment of inertia for a tube), the nerve mechanics should be negligible for any of the experimental nerve cuff materials. Given these calculations, this range shows that the PaC cuff minimises implant-tissue mechanical mismatch from a flexural rigidity standpoint, likely causing the near-absent FBR observed around the nerves for this case.

While the tested PE and PDMS cuffs were much thicker than PaC ones, PE and PDMS cuff dimensions were similar to those of thick-film fabricated nerve cuffs which typically employ these materials.

Clinically-used nerve stimulating cuffs have thicknesses in the millimetre range^7^. Although PDMS has worse insulating properties and is more difficult to fabricate into a thin film device than PaC, recent advances in brain and spinal cord implantable electrode arrays have employed PDMS with thicknesses of 100 – 200 μm ^27,28^. These lower thickness values would result in PDMS devices with lower mechanical mismatch to nerves, suggesting an alternative design to produce similarly long-term tissue-compatible cuffs.

### Long-term nerve recordings with ultraconformable cuffs

To examine the performance of the developed implants for long-term *in vivo* use, we performed forearm nerve recordings over 21 days in awake rats. An ultraconformable device implanted into the median, ulnar and radial nerves of the forelimb and with connections externalised through a headcap port was used to record nerve activity related to paw movement and sensation while the rat walked through a transparent tunnel (Fig. 3a). The cuffs were able to record action potential activity from nerves throughout the experiment period (Fig. 3b), with signal-to-noise ratios (SNR) and spike amplitudes remaining stable across the 21 days tested (Fig. 3c-d) (SNR across all timepoints: 9.7 ± 6.1, peak amplitude: 14.6 ± 6.6 µV, root mean squared noise: 3.1 ± 1.7 µV, mean ± standard deviation). No major change in microelectrode impedance or loss of functional electrodes was observed (Fig. 3e) indicating robust long-term implant performance. Spike sorting identified activity from multiple sources within the recorded traces of each of the three interfaced nerve with different temporal evolutions over the behavioural task (Fig. 3f, Supplementary Fig. 2), suggesting recordings from whole nerves could identify axon activity corresponding to different sensory and motor activity.

**Fig. 3.**
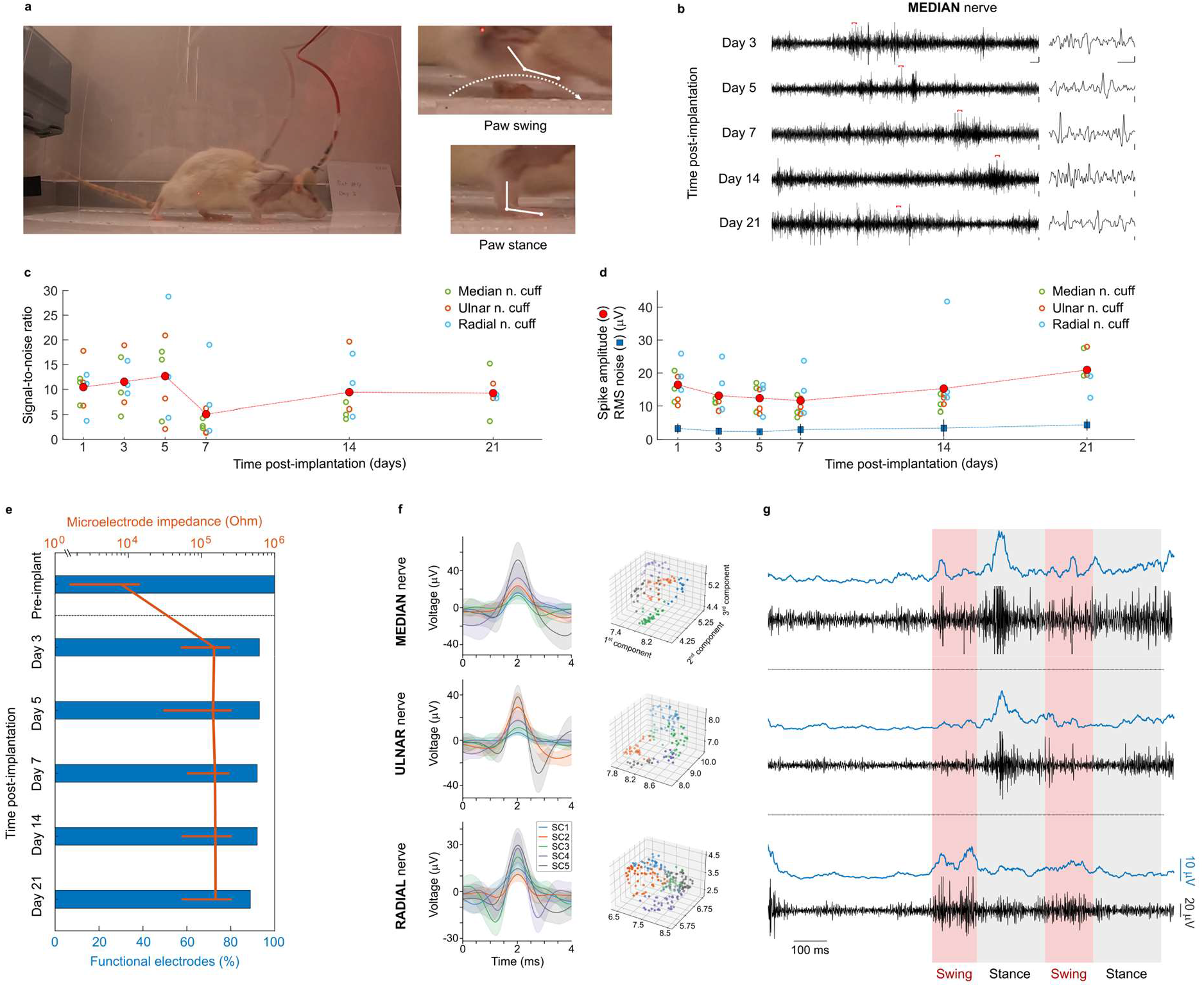
Ultraconformable nerve cuff arrays achieve stable recordings from their target nerves over chronic time periods. **a**, Photographs of behaviour task performed by animals during electrophysiology recordings. Animals are recorded while walking down a transparent tunnel. This allows the identification of nerve activity related to the swing and stance phases of walking of the implanted limb. **b**, Representative traces of recorded activity from the median cuff of a rat over a period of 21 days of implantation. Individual spikes can be seen throughout all timepoints (right, magnified from sections marked by red lines in traces on left). Scale bars: 100 ms and 10 µV (left) and 5 ms and 10 µV (right). **c-d**, Scatter plots of signal-to-noise ratio (SNR) (**c**) and spike amplitude (**d**) of recorded traces from median (green open circles), ulnar (orange open circles) and radial (cyan open circles) cuffs, showing stable quality of recording over the 21 day implantation period. Average of all cuffs represented by filled red circles. Root mean squared (RMS) of noise shown in (d) (mean: blue squares, standard deviation: black lines, for all cuffs) to represent spike detection threshold. *n* = 9 cuffs (6 for day 21) from 3 rats (2 for day 21). One rat had to be culled past the 14 day timepoint due to skin damage caused by the FFC of the implant. **e**, Microelectrode impedance and survival over the implantation period. Implantation leads to an impedance rise and the breakage of 7% of the microelectrodes, but these values remain thereafter stable. Functional microelectrodes are defined as having an impedance < 1 MOhm. *n =* 96 (Pre-implantation), 89 (Days 3-7), 88 (Day 14), 56 (Day 21) microelectrodes, from 3 rats (2 for day 21). **f-g**, Spike waveforms (**f**) and representative neural traces (black) and averaged RMS values (blue) (**g**) recorded from median, ulnar, and radial nerves simultaneously, 3 days post-implantation. Neural activity is seen to increase during the implanted limb stance phase of locomotion in median and ulnar nerves, and during the swing phase in radial nerve.

Examination of the relation between the pattern of recorded activity with animal behaviour identified periods of high nerve activity during walking. More specifically, high activity in median and ulnar nerves was seen to coincide with the stance portion of walking locomotion, while radial nerve activity coincided with the swing phase (Fig. 3g). This pattern of activity matched the innervation pattern of the three nerves, with median and ulnar nerves modulating paw flexion movements and sensation from the palm^17,18^ – expected to predominantly occur during stance. Radial nerve, instead, predominantly mediates the paw extension expected during the swing phase of walk^18^. Moreover, the distinct difference in recorded activity pattern across electrodes on different adjacent nerves supports that recorded activity is neural in origin and not originating from other sources such as electromyogram – a common contaminant of peripheral nerve recording.

### Sub-nerve resolution recording and signal velocity sorting

Implantable nerve cuffs are typically designed to record averaged activity from whole nerve^1,2,8^. By using low-impedance sub-mm microelectrode arrays that conform optimally to the nerve surface, we hypothesised that our devices would be able to record unique neural activity from portions of the nerve. In order to test this we compared the recordings of microelectrodes deployed as a ring around the same nerve (Fig. 4a).

**Fig. 4.**
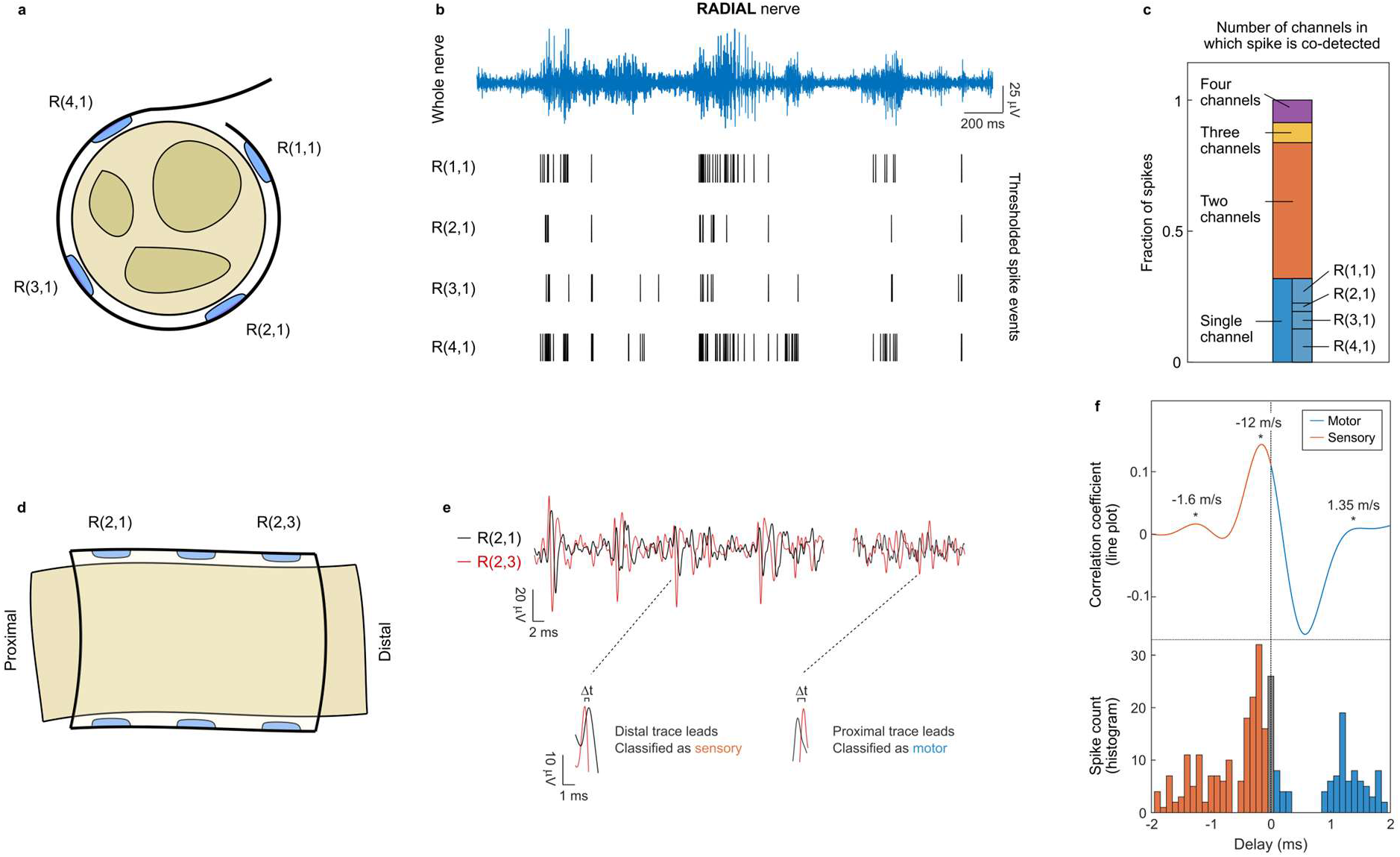
Ultraconformable cuff arrays achieve sub-nerve recording resolution without penetrating the nerve. **a**, Diagram of layout of one ring of microelectrodes around the circumference of the radial nerve. **b**, Spike events across four microelectrodes within a ring along the radial nerve circumference. The whole nerve recording is provided (blue) for reference. Spikes are detected in bursts in all microelectrodes, coinciding the swing phase during rat walking. Some spikes are detected across all microelectrodes, while others are detected uniquely in some of them. **c**, Quantification of spike coincidence across microelectrodes for activity shown in (b). Most neural spikes are detected across two microelectrodes simultaneously, or uniquely by a single microelectrode. All four microelectrodes record these unique spikes (breakdown of proportion represented in graph). Spikes are considered to be co-detected when they occur in several channels within 0.5 ms of each other. Co-detection is quantified by number of spikes, not by co-detection events (an event where four spikes are co-detected across all four microelectrodes is counted as four spikes). **d**, Diagram of layout of microelectrodes positioned along the length of the radial nerve. **e**, Neural activity recording from microelectrodes within two different rings in a cuff. Neural action potential spikes recorded across both rings exhibit a time delay, corresponding to their direction and velocity. This is highlighted at the bottom of the panel. **f**, Quantifications of radial nerve activity delay between microelectrodes within the most proximal and most distal rings in the array of the cuff over a 2.7 second awake recording. Top: quantification corresponding to the cross-correlation value between the nerve recordings of the two microelectrode at varying time delays. Bottom: histogram corresponding to the inter-spike interval between microelectrodes. Both analyses identify peaks of activity at delays corresponding to ∼1.6 m/s afferent, ∼12 m/s afferent, and ∼1.35 m/s efferent, calculated based on the 2 mm distance between microelectrode rings. Fast afferent/efferent activity is also detected at close to 0 ms delay.

In awake animals, all microelectrodes within the same ring recorded neural spikes in bursts which were synchronous across all microelectrodes (Fig. 4b, Supplementary Figs. 3-4). However, analysis of these spike bursts revealed that bursts were made up of different neural spikes across different microelectrodes. Coincidence analysis determined that most neural spikes were uniquely recorded by one or two microelectrodes within the ring, with few spikes being recorded across the entire ring of microelectrodes in a cuff (Fig. 4c, Supplementary Fig. 4). Moreover, all microelectrodes within a cuff ring could record unique spikes (Fig. 4c, Supplementary Fig. 4), indicating that spikes uniquely recorded by microelectrodes were not a result of poor performance of other microelectrodes in the ring. The median, ulnar and radial nerves contain two or three fascicles each, and each connect to synergist muscles and adjacent skin dermatomes^18,29^. This likely gives rise to the spike pattern observed - microelectrodes within the same ring record bursts of activity corresponding to the same overall movement or sensation, with each microelectrode recording neural spikes from different axons and fascicles. Our results therefore show that the developed ultraconformable cuffs are able to record selectively from portions of the nerve without the need to pierce the epineurium.

Having confirmed sub-nerve recording resolution in microelectrodes deployed around the circumference of the nerve, we examined recordings along the length of the nerve. We made use of the precise microelectrode arrangement enabled by microfabrication and the high recording quality shown by our ultraconformable cuffs to carry out velocity sorting of nerve signals in awake animals. Microelectrodes positioned on different locations along the nerve epineurium recorded the same spikes but with some amount of delay (Fig. 4d-e). We interpreted this delay as arising from the specific conduction velocity of the axon giving rise to that action potential. We observed both spikes which lagged in the microelectrode positioned distally along the nerve – efferent nerve signals – as well as spikes which lagged in the proximal microelectrode – afferent nerve signals (Fig. 4e). The abundance of spiking activity seen in awake nerve recordings and compact design of the cuff microelectrode arrangement made velocity analysis techniques previously developed and tested under anaesthesia^30–34^ unsuitable for our awake recordings. We developed two analysis strategies to determine conduction velocity, the first involving spike detection through thresholding via an adapted inter-spike histogram, and the second involving a cross-correlation calculation between the two recorded traces which utilised the entire recorded waveform. Both of these analytical strategies yielded agreeing results (Fig. 4f, Supplementary Fig. 5). We identified several populations of signal velocities corresponding to known sensorimotor nerve fibre types^35^, slow sensory 10 - 30 m/s (Aδ fibre velocity), slowest sensory 1 - 3 m/s (C fibre velocity) and slow motor 1 - 3 m/s (Aγ fibre velocity). The limited sampling rate used in typical electrophysiology recording hardware and the compact design required in chronically implanted devices limited sorting capabilities for fast sensorimotor fibres > 30 m/s (Aβ and Aα fibre velocities). However, we identified populations of spikes with almost no delay between microelectrodes, which likely corresponded to these fast fibres. Our results validated the neural velocity sorting capabilities in awake, physiological environments for the developed ultraconformable cuffs.

**Fig. 5.**
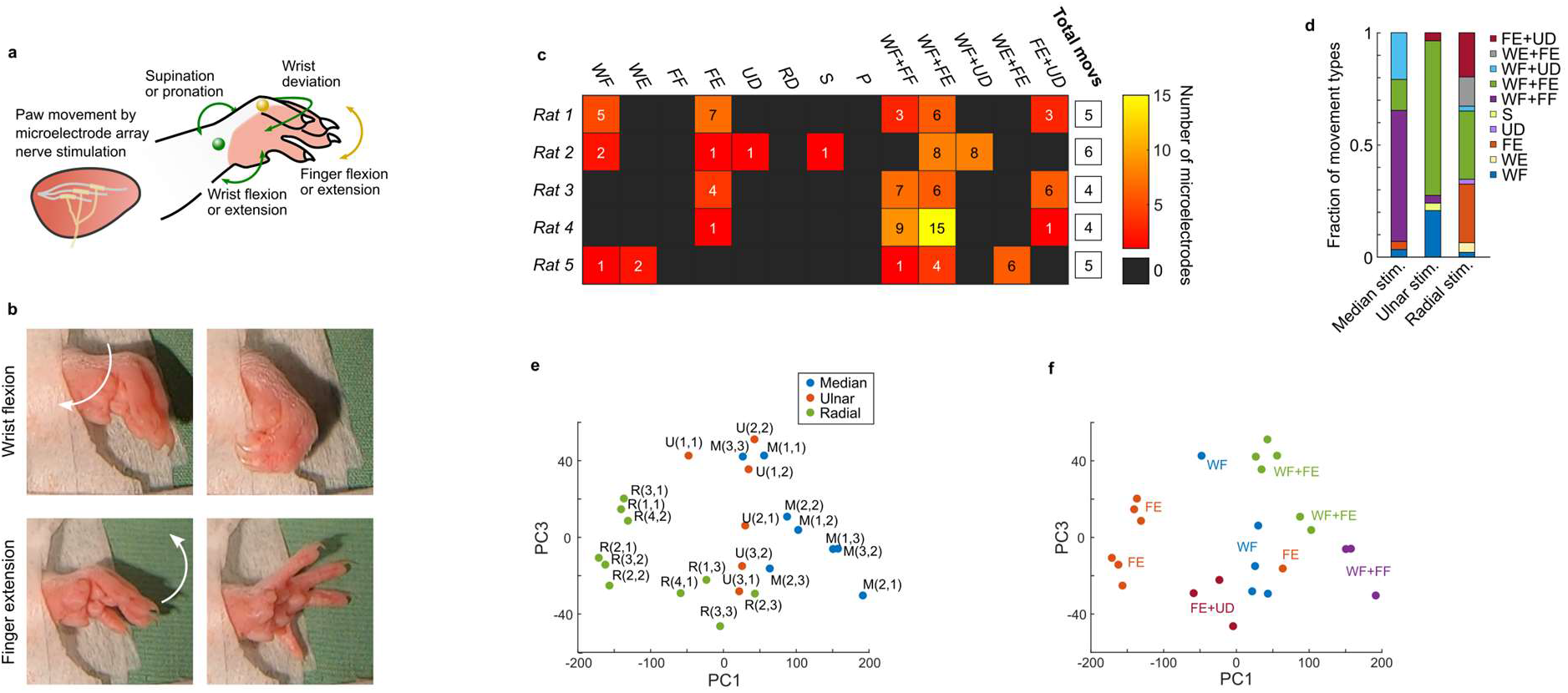
Ultraconformable cuff arrays achieve sub-nerve stimulation resolution. **a**, Diagram of the experimental setup. Stimulation of the three implanted nerves using individual microelectrodes causes movements of the wrist and fingers of the paw, in rats under anaesthesia. **b**, Pictures of examples of movements produced before (left) and during (right) nerve stimulation. **c**, Heatmap of quantification of movements produced by microelectrode stimulation across five rats. While stimulation on occasion caused simple movements, more often combination movements (containing “+” in name) were observed. Certain simple movements were never observed. Despite variations in movements across different implantations and animals, more than three movements were observed in every occasion, indicating higher than whole-nerve stimulation resolution. **d**, Quantification of the types of movements produced by stimulation using microelectrodes on the three different nerves. **e-f**, Plot of principal components of kinematic analysis of the various movements produced, for Rat 1. **e**, Movements are tagged based on the microelectrode that produced them. Clustering is seen across groups of microelectrodes within each cuff. **f**, Movements are tagged by the identified type of movement produced. Clustering indicates that many identified types of movements can be performed with different kinematics, indicating additional stimulation selectivity not captured in panel c). WF: wrist flexion, WE: wrist extension, FE: finger extension, UD: ulnar deviation, SD: sacral deviation, S: supination, P: pronation, WF+FF: wrist flexion and finger flexion, WF+FE: wrist flexion and finger extension, WF+UD: wrist flexion and ulnar deviation, WE+FE: wrist extension and finger extension, FE+UD: finger extension and ulnar deviation, Total movs: total number of different movements produced.

### Forelimb paw movement control using sub-nerve resolution stimulation

Implantable nerve interfaces are commonly utilised for neuromodulation applications through electrical stimulation. Similar to recordings, cuff interfaces are typically designed to stimulate whole nerves, while the more invasive penetrating interfaces selectively stimulate portions of the nerve^1,2,8^. We tested whether the developed ultraconformable cuffs could achieve selective stimulation of portions of the nerve, mirroring their recording capabilities, and comparable to the resolution achieved by penetrating devices.

We tested the neuromodulation capabilities of the cuffs by evaluating their ability to generate different forelimb paw movements (Fig. 5a-b). The median, ulnar and radial nerves are responsible for controlling hand movement, and nerve interfaces have been used in these nerves to restore hand function in amputation and spinal cord injury patients^15,19^. While rats have lower hand dexterity than primates, they are able to grasp objects with their forelimb paws. Testing in rats under anaesthesia showed that stimulation through the cuffs could achieve multiple different paw movements (Fig. 5c, Supplementary Fig. 6). While some variability in the range movements produced was observed across different implantations – likely due to small differences in nerve fascicle-microelectrode relative position – the cuffs could reliably produce movements including wrist and finger flexion as well as extension, which are key for hand function^15^. Although part of this movement selectivity was achieved through stimulation of the three different nerves, with flexion predominantly produced through median nerve stimulation and extension through radial nerve stimulation, stimulation of any one nerve using individual microelectrodes within the same cuffs could produce different movements (Fig. 5d). Overall, we observed more than three movements in all implantation experiments, with some implantations yielding up to six recognisable movements (Fig. 5c). These observations were indicative of the devices selectively stimulating portions of each nerve. We further confirmed this by performing an analysis of the movement kinematics, which showed that clusters of microelectrodes within the same nerve cuff could produce different movements (Fig. 5e-f, Supplementary Fig. 7).

As we could not explore activation characteristics of the forearm muscles via electromyogram recordings due to the small size of rats, we focused our analysis on the resulting hand movements. Our results confirmed that the developed implants were able of producing a variety of hand movements despite not penetrating into the nerve endoneurium. Moreover, while we only characterised movements produced by individual microelectrodes, we observed that these could be combined to produce more complex movements, either by simultaneous stimulation or activation in sequence (Supplementary Video 1). This supported that implants like the ones developed could be used for the restoration of hand movement in conditions such as spinal cord injury and stroke.

## Discussion

Existing nerve interfaces either use a high-invasive penetrating design to connect multi-electrode arrays with multiple portions of the nerve or use less invasive designs cuffing around whole nerves. Despite the long-term advantages of low-invasive designs, cuffs have traditionally been produced using rigid materials and bulky constructions^1,2,8^, resulting in generally low interface qualities and recording SNR. Stimulation with cuffs has been comparatively more successful, seeing use in human patients in the form of therapies such as vagus nerve stimulation^6,7^. However, clinical-use cuffs typically stimulate the entire nerve^7^, therefore limiting the scope for new applications, as most nerves contain multiple fascicles innervating different targets with physiological roles. While some studies have developed cuffs with electrode arrays designed to stimulate portions of the nerve, these have relied on the use of larger nerves with more limited therapeutic relevance where some degree of control is easier to achieve, such as the sciatic nerve to control leg muscles^36,37^ or slow deformation of larger nerves to spread out fascicles inside non-circular cuffs^38^.

The nerve cuffs developed in this work combined low-impedance microelectrode arrays with a low profile and ultraconformable design to achieve recording and stimulation sub-nerve resolutions previously unachieved by nerve cuff implants. From a recording perspective, the ultraconformable cuffs were able to record neural activity unique to certain points on the circumference of the nerve. The nerves of the brachial plexus to which the cuffs interfaced innervated synergist muscles and adjacent skin dermatomes, making interpretation of recorded sub-nerve neural activity difficult.

However, we observed that most neural spikes were recorded by either one or two of the three or four microelectrodes surrounding each nerve. As the median, ulnar and radial nerves in rats typically have two to four fascicles each^29^, it seems likely that the sub-nerve resolution recordings correspond to fascicle-specific activity. From a stimulation perspective, the ultraconformable cuffs could stimulate nerves of the forearm with sufficient resolution to not only generate different movements based on the cuff location of the microelectrode (indicative of sub-nerve stimulation resolution) but also produce physiologically meaningful movements such as wrist and finger extension (object release) and flexion (object grasp). Hand movement controlled via nerve stimulation is an active topic of research with applications in spinal cord injury and stroke. Recent advances using a penetrating neuromodulating interface (Transverse Intrafascicular Multichannel Electrode - TIME) have reproduced hand movements such as wrist and finger flexion and extension in non-human primates^15^. Albeit in rodents, the ultraconformable cuffs developed in this work were also capable of producing a comparable range of hand movements despite their less invasive design.

The extraction of conduction velocity and direction from nerve recordings from awake freely-moving animals is a novel feature implemented in our ultraconformable cuffs. Action potential conduction through axons occurs at velocities ranging from approximately 0.4 to 120 m/s, with different types of axon fibres having different conduction velocities. As different nerve fibre populations are associated with different physiological and pathological roles^39^, a method to selectively identify the activity of specific velocities is a highly valuable research and clinical tool. Methods to sort recorded nerve activity based on conduction velocity and direction have long existed^30^ and have mostly been used and validated in animals under anaesthesia to analyse the velocity of electrically-evoked compound action potentials^32,33^. However, the lack of implantable devices with high quality recordings and precise electrode positioning has prevented their translation to more physiologically-relevant awake models. The high interface quality and compact design of the cuffs developed in this work allowed us to implement this feature. Our devices recorded action potential conduction velocities in the expected physiological range. Earlier velocity analysis methods often relied on the addition of individual action potentials with varying amounts of delay, producing as a readout the maximum amplitude of the *delay-and-add* trace^30,31,33^. This method was not well suited for velocity sorting of large volumes of bidirectionally transmitted spikes from awake animals, leading us to develop two new analytical methods. While the nerves targeted and behavioural setup used in this work did not allow us to manipulate the activity of select nerve fibre populations, the similar results yielded by both analysis methods and agreement with known fibre population velocities^35^ supports the validity of the velocity sorting capabilities of our cuffs.

One of the main challenges of peripheral nerve interfaces is chronic stability. Unlike the CNS, nerves are located in highly dynamic environments characterised by large movements and forces and are not encased in protective bone structures. These properties can make breakages in implants a common occurrence and exacerbate the tissue damage and FBR generated due to tissue-implant mechanical mismatch^8^. The use of robust thick-film fabricated designs has enabled whole-nerve stimulation in cuffs lasting years^40^ and the more fragile, microfabricated penetrating nerve interfaces have also been used for stimulation in humans for periods of months^41^. Long-term recordings are much more difficult to achieve as they heavily rely on the presence of a high quality tissue-electrode interface, which can quickly degrade due to chronic tissue inflammation and FBR. Traditional cuff electrodes generally perform poorly in nerve recordings due to their overall worse nerve-electrode interface, with few applications making use of them outside of animal models under acute anaesthesia^2,8,33,34,42^. Penetrating devices perform better. Slanted Utah electrode arrays have been used to record nerve activity in humans for up to 27 days, albeit with some progressive decrease in the number action potential-recording electrodes^43^ and with the development of chronic inflammation, which could pose a challenge to long term use^44^. Our ultraconformable cuffs achieved stable recordings in awake animals for the 21 days over which we performed recordings, with minimal loss of electrodes, negligible change in impedance after implantation and little formation of fibrotic scar tissue. Moreover, the cuff design provides a number of advantages compared to penetrating devices with regards to maintaining a stable recording interface. The proximity of penetrating devices to axons makes them more sensitive to the build-up of fibrotic tissue at the interface, in contrast to cuffs designed to record through a ∼50 μm fibrous epineurium. Secondly, cuffs transfer forces through the epineurium – a collagen-rich tissue responsible for protecting the nerve – whereas penetrating devices interact with and cause chronic inflammation in the much more fragile endoneurium where axons are located.

Overall, the developed ultraconformable nerve cuffs serve as a tool for a wide range of applications. The ability to record chronically while identifying active nerve fibre populations in awake animals provides a new avenue to study the physiology of nerves and innervated structures, particularly when combined with fascicle-specific recording and neuromodulation enabled by this technology. Clinically, the developed technology may not only see use in the control of hand movement but may also serve as a less invasive and more chronically-stable neuromodulation tool than penetrating devices, potentially also enabling the clinical use of nerve recordings for diagnostic or closed-loop applications.

## Methods

### Device fabrication

Implants were fabricated using photolithographic techniques for flexible electronic devices^12^. PaC, from dichloro-p-cyclophane (Specialty Coating Systems, Indianapolis, IN, USA) was deposited (Specialty Coating Systems Labcoater – Specialty Coating Systems, Indianapolis, IN, USA) at a 2 μm thickness onto a Si wafer. Gold tracks were patterned (MA/BA6 or MJB4 mask aligner – Süss Microtec, Garching, Germany) using AZ 5214E photoresist and AZ 400K developer or using AZ nLOF 2035 photoresist with AZ 726 MIF developer (Microchemicals GmbH, Ulm, Germany). A 10 nm Ti adhesion layer followed by a 100 nm Au layer was deposited (Lesker e-beam Evaporator – Kurt J. Lesker Company, Jefferson Hills, PA, USA) onto the PaC layer, and a gold lift-off was performed using acetone. A second layer of PaC was deposited, insulating the gold tracks. The outline of the device was patterned using AZ 10XT 520cP (Microchemicals GmbH, Ulm, Germany) photoresist with AZ 726 MIF developer to create an etch mask. The outline was etched using reactive ion etching (PlasmaPro 80 Reactive Ion Etcher (RIE) – Oxford Instrument, Abingdon, UK), using 8 sccm CF_4_, 2 sccm SF_6,_ and 50 sccm O_2_ at 60 mTorr. Miro-90 Concentrated Cleaning Solution (Cole-Parmer, Vernon Hills, IL, USA) was spin-coated onto the devices as an anti-adhesive layer before a third layer of 2 μm of PaC was deposited onto the wafer. The electrodes and connector pads were patterned and etched in the same manner as the device outline. Electrodes were then spin-coated with a PEDOT:PSS coating, consisting of 5 v/v% ethylene glycol, ∼30 µL of DBSA, 1 v/v% GOPS (Sigma Aldrich, St. Louis, MI, USA), remainder PEDOT:PSS (Clevios PH 1000 – Heraeus, Hanau, Germany). After coating, the anti-adhesive PaC layer was peeled off, leaving discretely-coated electrodes. Finally, implants were removed from the Si wafer, and an FFC (Mouser Electronics, Mansfield, TX, USA) was bonded (Finetech Bonder Fineplacer® System – Finetech GmbH, Berlin, Germany) onto the connector using anisotropically conductive film (5 μm particulate) (3TFrontiers, Singapore) to create the implant.

### Surgical implantation

All animal procedures were carried out in accordance with the UK Animals (Scientific Procedures) Act, 1986. Work was approved by the Animal Welfare and Ethical Review Body of the University of Cambridge, and was approved by the UK Home Office (project licence numbers PFF2068BC and PP5478947). Experiments were conducted on Sprague Dawley (nerve stimulation) or Lewis (chronic nerve electrophysiology recordings) female rats ∼150-200 g in weight (Charles River, UK). Rats were group-housed in individually ventilated cages with ad libitum access to food and water for the duration of the study.

Surgical implantation of devices was carried out under isoflurane anaesthesia (2.25% v/v in medical oxygen). Body temperature was monitored and maintained using a thermal blanket. For non-recovery (nerve stimulation) experiments, an incision was made in the ventral portion of the arm. The bundle of radial, ulnar and median nerves was accessed between the humerus and the triceps muscle at the upper-arm level. The three cuffs of the ultraconformable device were implanted by wrapping around each of the three nerves, and sealed shut using a drop of Kwik-Cast silicone (World Precision Instruments) applied between the locking tab and the PaC connector. Care was taken not to allow silicone to flow into or around the cuff portion of the implant.

For chronic electrophysiology recording experiments, the nerves were accessed at a similar height dorsally, and cuffs were similarly implanted. The PaC and FFC connectors were routed subcutaneously to the head of the animal, and held in place using a 6-0 suture (Ethilon Nylon, Ethicon). The PaC portion of the connector was reinforced with a thin layer of Kwik-Cast, applied after cuff implantation. The connector was routed to a custom 3D-printed headcap, which was secured to the head of the animal using surgical cement (Unifast Trad, Billericay Dental Supply Co Ltd; Super Bond C&B, Prestige Dental Products Ltd) and drilled steel screws (M0.8, US Micro Screw). An additional screw was drilled above the cerebellum and connected to using a steel wire to use as an electrophysiology ground. The FFC and ground wire were connected to a custom PCB integrated into the headcap, which provided the connections to perform electrophysiology in awake animals.

For the chronic FBR immunohistochemistry study, the radial nerve was accessed ventrally at the same height, and a cuff consisting of either a medical-grade PDMS tube (Syndev, 0.63 mm inner diameter, 1.19 mm outer diameter, 228-0254 VWR), implantable cannula PE tube (0.6 mm inner diameter, 1.1 mm outer diameter, 504280, World Precision Instruments), or a PaC cuff was implanted. The tube implants were produced by slicing open a 2 mm-long portion of tube and allowing it to wrap around the nerve. The PaC cuffs consisted of median or radial cuffs from ultraconformable cuff devices, cut at the PaC connector portion of the device. These were implanted by wrapping around the nerve and sealed with a drop of Kwik-Cast silicone between locking tab and PaC connector portion of the implant. Care was taken to not allow silicone to flow into or around the cuff portion of the implant.

In chronic implantation surgeries (FBR and electrophysiology recordings), incisions in the arm were sutured closed (5-0 Monocryl sutures, Ethicon) and animals were recovered from anaesthesia. Anaelgesia was provided for two days following surgery (Metacam, oral suspension), and were kept under post-operative observation for three days following surgery.

### Immunohistochemistry

Four weeks post-implantation animals were humanely killed by rising CO_2_ concentration. The implanted nerves were accessed and dissected out together with their implant, and fixed in formalin overnight at 4ºC. The cuffs were then carefully removed under a dissection microscope and the nerves trimmed to keep only the portion which had been surrounded by a cuff. The nerve samples were then transferred into phosphate buffered saline with 30% w/w sucrose for cryoprotection, mounted in optimal cutting temperature (OCT) compound (361603E, VWR), and sectioned in a cryostat (CM1900, Leica) into 10 µm-thick cross-sections of the nerves.

For staining, sections were blocked in 5% donkey serum (D9663, Sigma) in phosphate buffered saline and 0.1% sodium azide for 1 hour at room temperature. Anti-αSMA (1/100 dilution, ab7817, Abcam) and Anti-β3 tubulin (1/1000 dilution, ab18207, Abcam) antibodies in blocking medium were added for 3 hours at room temperature in the dark. This was followed by incubation in secondary antibodies (Donkey anti-Rabbit IgG (H+L) Highly Cross-Adsorbed Secondary Antibody, Alexa Fluor 555; and anti-Mouse, Alexa Fluor 488; Invitrogen) for 3 hours at room temperature in the dark. Finally, sections were incubated in Vector TrueVIEW Autofluorescence Quenching Kit (Vector Laboratories) for 3 minutes at room temperature, mounted, and imaged in a Axioscan Slide Scanner (Zeiss). Between incubation steps, sections were washed three times, 10 minutes per wash, with blocking medium.

Analysis was carried out using custom scripts in MATLAB (Mathworks, 2016b) and Fiji (v1.48, National Institutes of Health, USA) in 20X magnification images. These produced a stain intensity profile of αSMA as a function of depth into the nerve, starting from the outermost portion of the circumference where the implant was located as defined by the user. Intensity was normalised into a ratio relative to αSMA intensity in the middle of the nerve cross-section of each nerve. Statistical analysis and data plotting was carried out in MATLAB (Mathworks, R2016b).

### Tensile tests

For mechanical testing of PaC, a PaC film was deposited onto a Si wafer as above at 6 μm thickness. PaC films were cut (Silhouette Portrait – Silhouette America, Inc., Lindon, UT, USA) into dog-bone mechanical testing samples. Samples were tested to failure (Tinius Olsen 1 ST – Tinius Olsen, Horsham, PA, USA) at a rate of 10 mm/min. Stress and strain were tabulated and used to calculate the elastic modulus. For mechanical testing of PE and PDMS, tubes of each material were tested as is within the elastic regime at a rate of 0.8 mm/min. Stress and strain were tabulated and used to calculate the elastic modulus.

### Nerve awake electrophysiology recordings

Electrophysiology recordings were carried out from rats at various timepoints post-implantation. Recordings were performed through an Intan RHS stim/recording system (Intan Technologies) connected to the PCB on the rat headcaps, amplified (x 192), bandpass filtered between 1 Hz and 7.5 kHz and sampled at a rate of 30 kHz. The cerebellar screw connected to this PCB was used as a ground for these recordings. Impedance measurements were performed using this same system and setup. Recordings were performed while rats walked through a transparent acrylic tunnel with an open top (500 mm long by 80 mm wide). Animals were video recorded during these tasks using a high frame-rate camera (GoPro Hero 6 Black, 120 fps). The walking task was repeated five times per rat per timepoint. The electrophysiology and video recordings were synchronised using an LED driven by the electrophysiology system. Swing and stance portions of the rat movement were identified manually from the recorded videos. Recording experiments were performed over 21 days post-implantation. One rat had to be culled 14 days post-implantation due to skin damage caused by the device FFC connector.

### Electrophysiology recording analysis

Electrophysiology analysis and data plotting was performed in MATLAB (Mathworks, R2016b) and Python. Recorded traces were imported and bandpass filtered using a 4^th^ order Butterworth filter with cutoff frequencies 300 and 2000 Hz. For each microelectrode, recordings were converted into a bipolar configuration through signal subtraction where the electrode used as a reference depended on the analysis being carried out. For whole nerve analysis (Fig. 3), the whole nerve electrode from the corresponding cuff of each microelectrode was used. For waveform sorting, neural events (‘spikes’) used as input to the spike sorting algorithms were identified using an adaptive threshold based on the smallest of constant false-alarm rate (SO CFAR) filter^45,46^, with the parameters (window duration of 150 ms on each side, guard cell duration of 10 ms on each side, and threshold level of 3 standard deviations from the mean) being heuristically chosen. The extracted waveforms were initially transformed to reduce dimensionality using the uniform manifold approximation and projection (UMAP) method. The new 3D space corresponding to the waveforms was used as input to an unsupervised clustering algorithm. The K-Means algorithm was used to keep consistency among trials on the number of identified clusters (heuristically set to 5 after preliminary analysis). Finally, a sliding window of 1 second was used to compute the spike rate of the clusters within each window, and the evolution of the metric over time was plotted (see Supplementary Fig. 2). SNR from recordings was calculated as the ratio between the variances of a randomly chosen burst of activity and of a period of no activity (noise), for each of the three cuffs in each implant at every timepoint. Using these same activity and baseline periods, root mean squared values for baseline and mean spike amplitude for activity were calculated (spikes detected using a threshold of 2.5 times the noise root mean squared).

In sub-nerve resolution (Fig. 4) nerve recording selectivity analysis, bipolar referencing was carried out between each microelectrode and the average of the other microelectrodes within that same ring. Microelectrodes within the same column in the array were deployed as a ring around the cuffed nerve when implanted. Co-detection analysis was carried out by first performing spike detection over all the referenced recordings from microelectrodes within the same ring (detection of peaks above 6.5 µV and with a minimum distance of 0.5 ms), and second by analysing whether each identified spike had an equivalent spike in any other channel within that same ring. A spike was considered to be co-detected when the peak of the spike is within 0.5 ms across two or more channels. Depending on whether the spike was co-detected across four (in case of the radial nerve cuff), three, two microelectrodes, or uniquely recorded in one microelectrode, it was categorised as “four channels”, “three channels”, “two channels”, or “single channel”. This analysis was performed for every spike in each microelectrode recording, meaning that for example a spike co-detected across three channels was counted three times. Single channel-recorded spikes were also categorised by the microelectrode within the ring from which they were recorded.

Spike delay analysis was carried out by comparing recorded signals from two bipolar-referenced microelectrodes from rings 2 mm apart within the array. Analysis was carried out in two ways, both intended for fast processing of large volumes of recorded action potential data. The first implementing a thresholding strategy and the second utilising the entire recorded trace. The first approach implemented a thresholding strategy. Spike detection was carried out over the traces of the two microelectrodes (detection of peaks above 6.5 µV and with a minimum distance of 0.5 ms). For each identified spike in a microelectrode, the spike closest in time in the other microelectrode recording was found and the delay between the two was calculated. The analysis was performed for spikes over both microelectrodes. Spike delays between -2 and +2 ms were grouped into a histogram with bin size 0.1 ms. The second approach made use of the entire recorded trace to avoid any velocity data distortion introduced by thresholding. The correlation coefficient was calculated between the two microelectrode traces when delays ranging from -2 to +2 ms were introduced. Results from the two analyses were plotted, and delays in spike/correlation were converted to conduction velocity and direction of neural activity using the distance and position on the nerve of the two microelectrodes.

### Nerve stimulation

Nerve stimulation was carried out on animals under isoflurane anaesthesia (1.75% v/v in medical oxygen) using an RHS stim/recording system (Intan Technologies). The system was connected to the implanted nerve cuffs with 32 microelectrodes, and used a larger electrode present in all three cuffs as a ground. Nerve stimulation was carried out through individual microelectrodes via trains of 100 biphasic pulses (100 µs and 100 µA per phase, 10 ms period). The resulting muscle contractions and movements of the implanted paw were recorded using a high frame rate camera (GoPro Hero 6 Black, 120 fps) from two angles (top and side of paw).

Analysis of movements and movement kinematics as well as data plotting was carried out in MATLAB (Mathworks, R2016b). Movements were classified by hand blindly (experimenter blinded to what movement corresponds stimulation through which microelectrode) based on the closest match between a series of predefined movements. These predefined movements were finger extension or flexion, wrist extension or flexion, radial or ulnar deviation, and paw pronation or supination. Recorded movements were identified as producing either a single of these predefined movements, a combination of two of these movements, or no movement at all. For kinematic analysis, video recordings were imported and position markers were placed over the paw for the movement produced by each microelectrode stimulation, as well as the paw resting position. Markers were placed on the tip of the middle digit (D3) and middle knuckle (top view video) and tip of thumb digit (D1), middle digit (D3) and little digit (D5) (side view video). X and Y positions of the markers were normalised to additional markers used as position origin on the base of the wrist (both top and side view). The position kinematics were processed by principal component analysis, and plots of the first and second, as well as first and third principal components, were produced.

## Data availability

The main data supporting the results in this study is available within it and its supplementary information. Source data for the figures in this study are available on Figshare with the identifier https://doi.org/10.6084/m9.figshare.21892986 (ref. ^47^). The raw datasets generated during the study are too large to be publicly shared, but are available for research purposes from the corresponding author on reasonable request.

## Code availability

Custom scripts used to carry out stain, electrophysiology and kinematic analysis are available in the following repository: https://doi.org/10.5281/zenodo.7544535.

## Acknowledgements

The authors would like to acknowledge Dr. Rachana Acharya and the Mechanical Testing Laboratory of the Department of Materials Science & Metallurgy of the University of Cambridge for their support in the tensile test experiments. A.C.L. acknowledges support from the University of Cambridge for a Borysiewicz Interdisciplinary Fellowship and the Wellcome Trust for a Junior Interdisciplinary Fellowship. A.J.B. acknowledges support from his Cross-disciplinary Fellowship (LT000034/2020-C) from the Human Frontier Science Program (HFSP) Organization. A.G.G. acknowledges funding support from the Royal Commission for the Exhibition of 1851. J. G. acknowledges the support from the German Research Foundation (Deutsche Forschungsgemeinschaft DFG via Research Fellowships Gz. GU 2073/1-1). S.V.B. acknowledges support from the W.D. Armstrong Studentship. This work was funded by the ECH2020 FUTURE & EMERGING TECHNOLOGIES (FET) projects BrainCom (732032) and MITICS (964677).

## Author contributions

A.C.L. designed the implants, carried out all implantation surgeries and electrophysiology experiments. A.J.B. carried out all fabrication of devices, tensile test experiments and mechanical analysis. A.C.L. and A.G. performed analysis of electrophysiology data. A.C.L., J.G. and S.V.B. designed and fabricated the awake electrophysiology connections and access ports. S.H. performed immunohistochemistry experiments. A.C.L., D.G.B. and G.G.M. designed the experimental plan. A.C.L. wrote the manuscript. All authors contributed to the review and revision of the manuscript.

