## Supplementary Information for "Ultraconformable cuff implants for long-term bidirectional interfacing of peripheral nerves at sub-nerve resolutions"

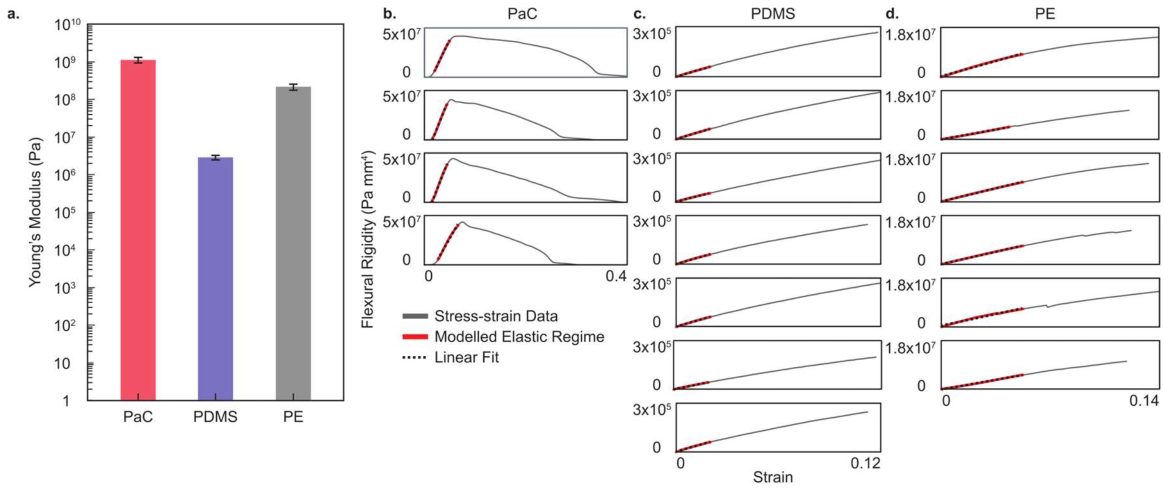


**e.** ***Moment of Inertia:***

***Moment of Inertia for a tube:***

$$I = {}^{J_0}/_{2}\,; I = \text{Moment of Inertia}, J_0 = \text{Polar Moment of Inertia}$$

$$J_0 = \int_A r^2\, dA;\, dA = 2\pi\rho d\rho\ (\text{for a circle})$$

$$J_0 = \int_{r_i}^{r_o} \rho^2 2\pi\rho d\rho$$

$$J_0 = \frac{2\pi\rho^4}{4}\Big|_{\rho = r_i}^{r_o}$$

$$J_0 = \frac{\pi}{2}(r_o^4 - r_i^4)$$

$$I = \frac{\pi}{4}(r_o^4 - r_i^4)$$

***Moment of Inertia for a flat plate:***

$$I = \int_A y^2 dA\,;\, dA = dxdy$$

$$I = \int_0^h \int_0^b y^2 dxdy$$

$$I = \int_0^h y^2 b dy$$

$$I = \frac{by^3}{3}\Big|_{y = 0}^{h}$$

$$I = \frac{bh^3}{3}$$

***Moment of Inertia for a cylinder:***

$$I = {}^{J_0}/_{2}\,; I = \text{Moment of Inertia}, J_0 = \text{Polar Moment of Inertia}$$

$$J_0 = \int_A r^2\, dA;\, dA = 2\pi\rho d\rho$$

$$J_0 = \int_{r_i}^{r_o} \rho^2 2\pi\rho d\rho$$

$$J_0 = \frac{2\pi\rho^4}{4}\Big|_{\rho = 0}^{r}$$

$$J_0 = \frac{\pi}{2} r^4$$

$$I = \frac{\pi}{4} r^4$$

***Parameters:***

***PaC:***
$E = 1.13 \times 10^9\ Pa$
$r_o = 0.3\ mm$
$r_i = 0.296\ mm$
$h = 0.004\ mm$

***PDMS:***
$E = 2.86 \times 10^6\ Pa$
$r_o = 0.595\ mm$
$r_i = 0.315\ mm$
$h = 0.28\ mm$

***PE:***
$E = 2.17 \times 10^8\ Pa$
$r_o = 0.55\ mm$
$r_i = 0.3\ mm$
$h = 0.25\ mm$

***Nerve:***
$E \approx 60\ Pa$
$r = 0.3\ mm$

***Flexural Rigidity (EI):***

***PaC:***
$I_{tube} = 3.33 \times 10^{-4}\ mm^4$
$I_{plate} = 2.13 \times 10^{-9}\ mm^4$
$EI_{tube} = 3.76 \times 10^5\ Pa\ mm^4$
$EI_{plate} = 2.41\ Pa\ mm^4$

***PDMS:***
$I_{tube} = 9.07 \times 10^{-2}\ mm^4$
$I_{plate} = 7.32 \times 10^{-4}\ mm^4$
$EI_{tube} = 2.59 \times 10^5\ Pa\ mm^4$
$EI_{plate} = 2.09 \times 10^3\ Pa\ mm^4$

***PE:***
$I_{tube} = 6.55 \times 10^{-2}\ mm^4$
$I_{plate} = 5.21 \times 10^{-4}\ mm^4$
$EI_{tube} = 1.42 \times 10^7\ Pa\ mm^4$
$EI_{plate} = 1.13 \times 10^5\ Pa\ mm^4$

***Nerve:***
$I_{cylinder} = 6.36 \times 10^{-3}\ mm^4$
$EI_{cylinder} = 0.38\ Pa\ mm^4$

**Supplementary Fig. 1 | Mechanical testing and flexural rigidity calculations different nerve cuff materials and nerve. a,** Young's modulus distribution for each material. **b-d,** testing curves for (**b**) PaC, (**c**) PDMS and (**d**) PE. **e,** Flexural rigidity calculations.

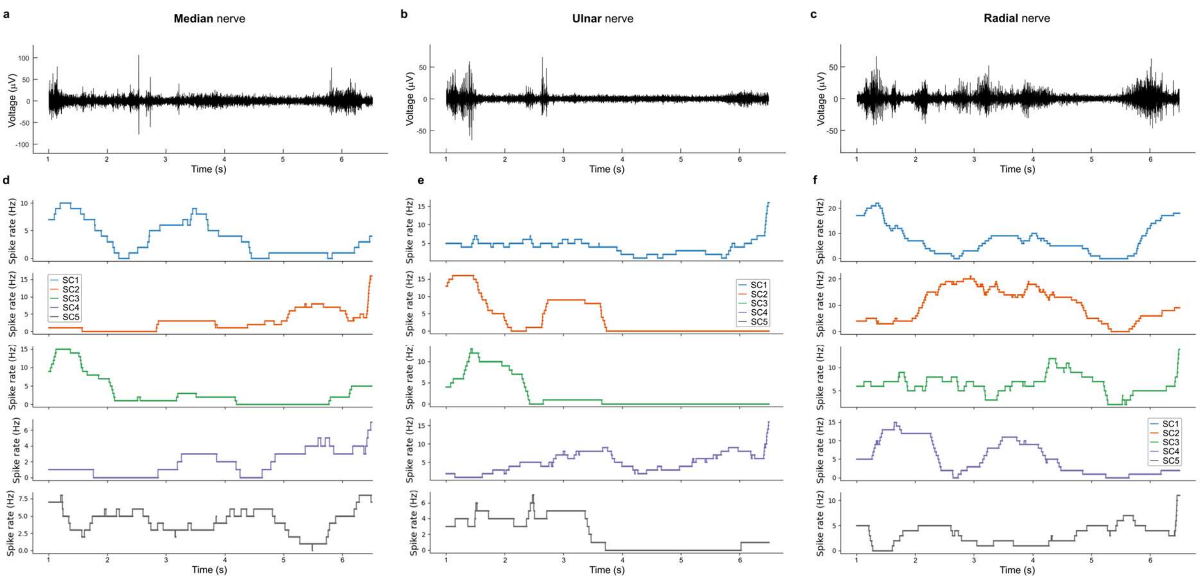


**Supplementary Fig. 2 | Temporal evolution of spike clusters among the three nerve cuffs. a-c**, Simultaneous neural recordings from median (**a**), ulnar (**b**) and radial nerves. **d-f**, Spike rates for five spike clusters sorted from recordings in (a-c), with a one second averaging window. Different clusters contribute predominantly to certain activity bursts in whole nerve recording traces. Traces and clusters obtained from an awake rat 3 days post-implantation (same as Fig. 3f-g).

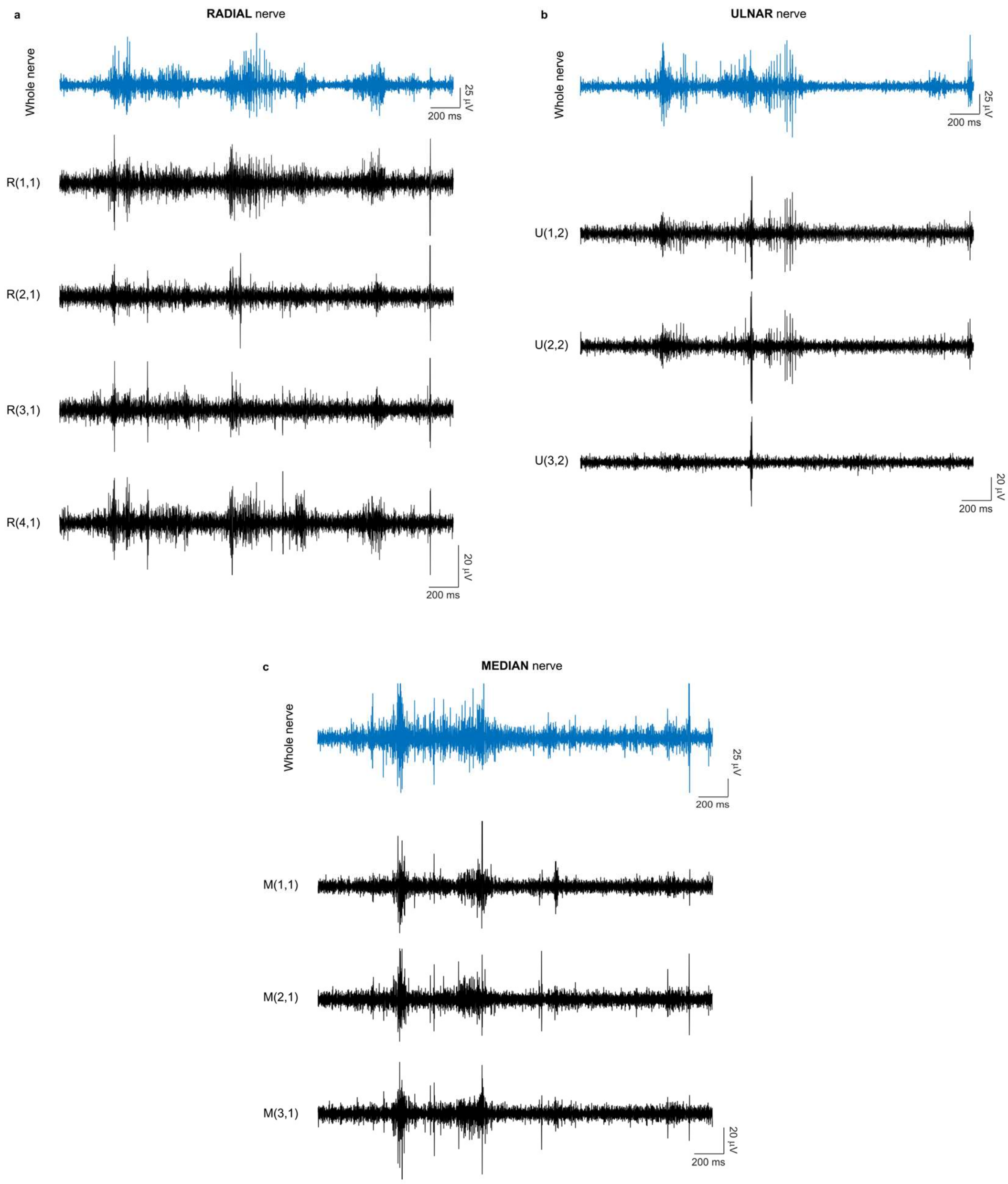


**Supplementary Fig. 3 | Cuff recordings along the circumference of radial, ulnar and median nerves.** **a**, Recording from whole nerve electrode (top, blue) and from four microelectrodes in a ring along the circumference (bottom, black) of the radial nerve. Microelectrode traces correspond to those on which event thresholding is carried out in Figure 4b. **b-c**, Recordings from whole nerve and ring of microelectrodes from ulnar (**b**) and median (**c**) nerves. All three nerve recordings are obtained simultaneously from the same rat, 3 days post-implantation.

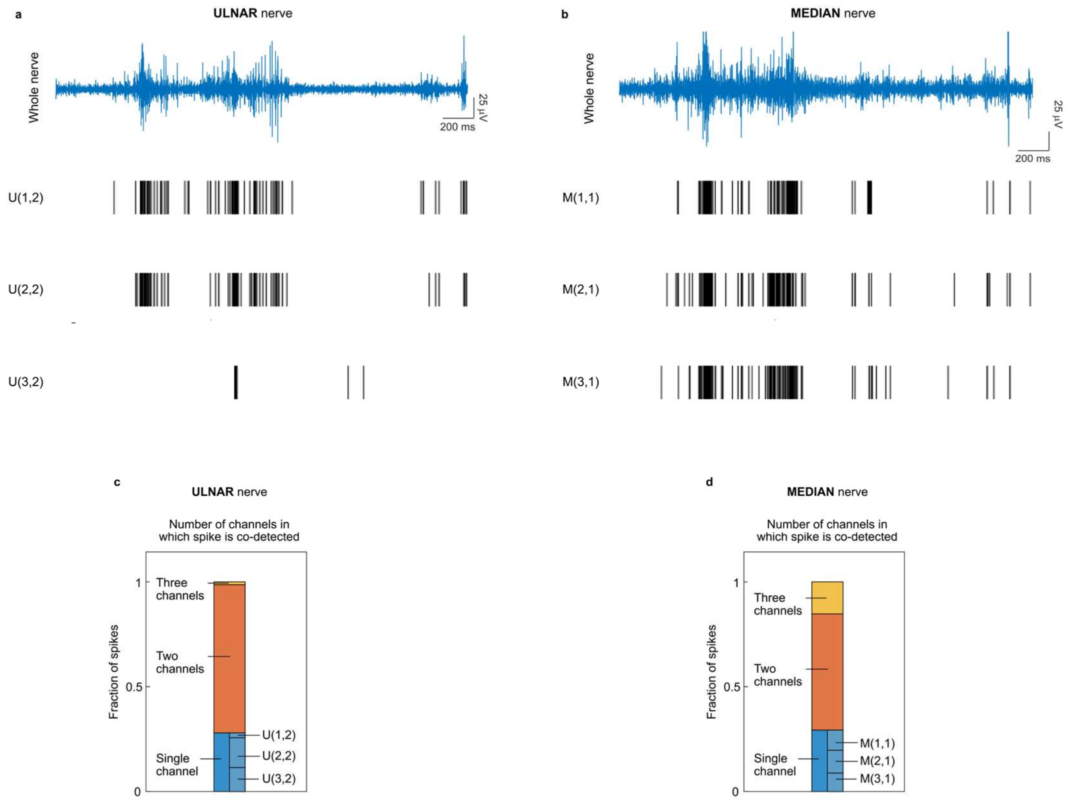


**Supplementary Fig. 4 | Circumference sub-nerve recording capabilities of ultraconformable cuff microelectrodes in ulnar and median nerves. a-b**, Spike events across three microelectrodes within a ring along the ulnar (**a**) and median (**b**) nerve circumference. The whole nerve recording is provided (blue) for reference. Spike thresholding is carried out over recording traces from Supplementary Figure 2b-c. **c-d**, Quantification of spike coincidence across microelectrodes for activity shown in (a-b). Recordings and coincidence calculation obtained simultaneously from those of Figure 4b-c from an awake rat 3 days post-implantation.

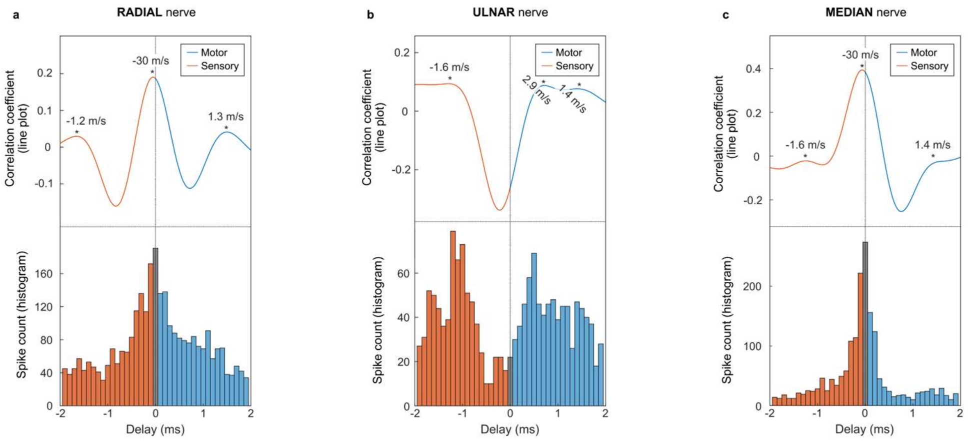


**Supplementary Fig. 5 | Nerve activity direction and velocity analysis using longitudinal sub-nerve recording capabilities of ultraconformable cuff microelectrodes. a-c**, Quantifications of nerve activity delay between microelectrodes within the most proximal and most distal rings in the array of each cuff, for radial (**a**), ulnar (**b**) and median (**c**) nerves. Top: quantification corresponding to the cross-correlation value between the nerve recordings of the two microelectrodes at varying time delays. Bottom: histogram corresponding to the inter-spike interval between microelectrodes. Analysis identifies different peaks of sensory and motor activity at various values, differing across the three nerves. Analysis performed over a 15-long recording obtained simultaneously from all three nerves from an awake animal 3 days post-implantation. Recordings used for analysis differ from those used in Figure 4f. Velocities are calculated based on the 2 mm distance between microelectrodes.

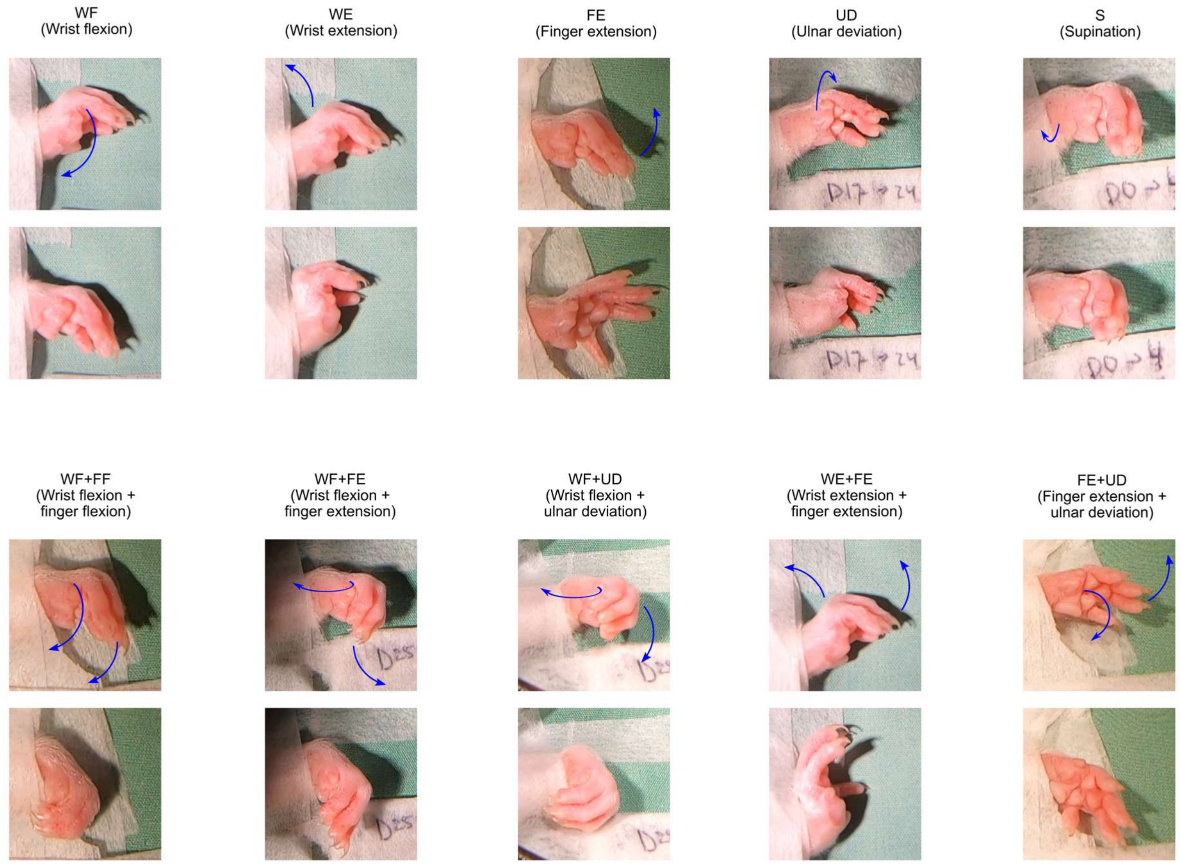


**Supplementary Fig. 6 | Picture gallery of paw movements induced by nerve stimulation through ultraconformable cuffs.** Pictures before (top) and during (bottom) nerve stimulation. Blue arrow indicates direction of movement. Composite movements (bottom row, indicated by "+" in their name) consist of two simultaneous movements. Movements can occur around both wrist (WF, WE, UD, S) or fingers (FF, FE).

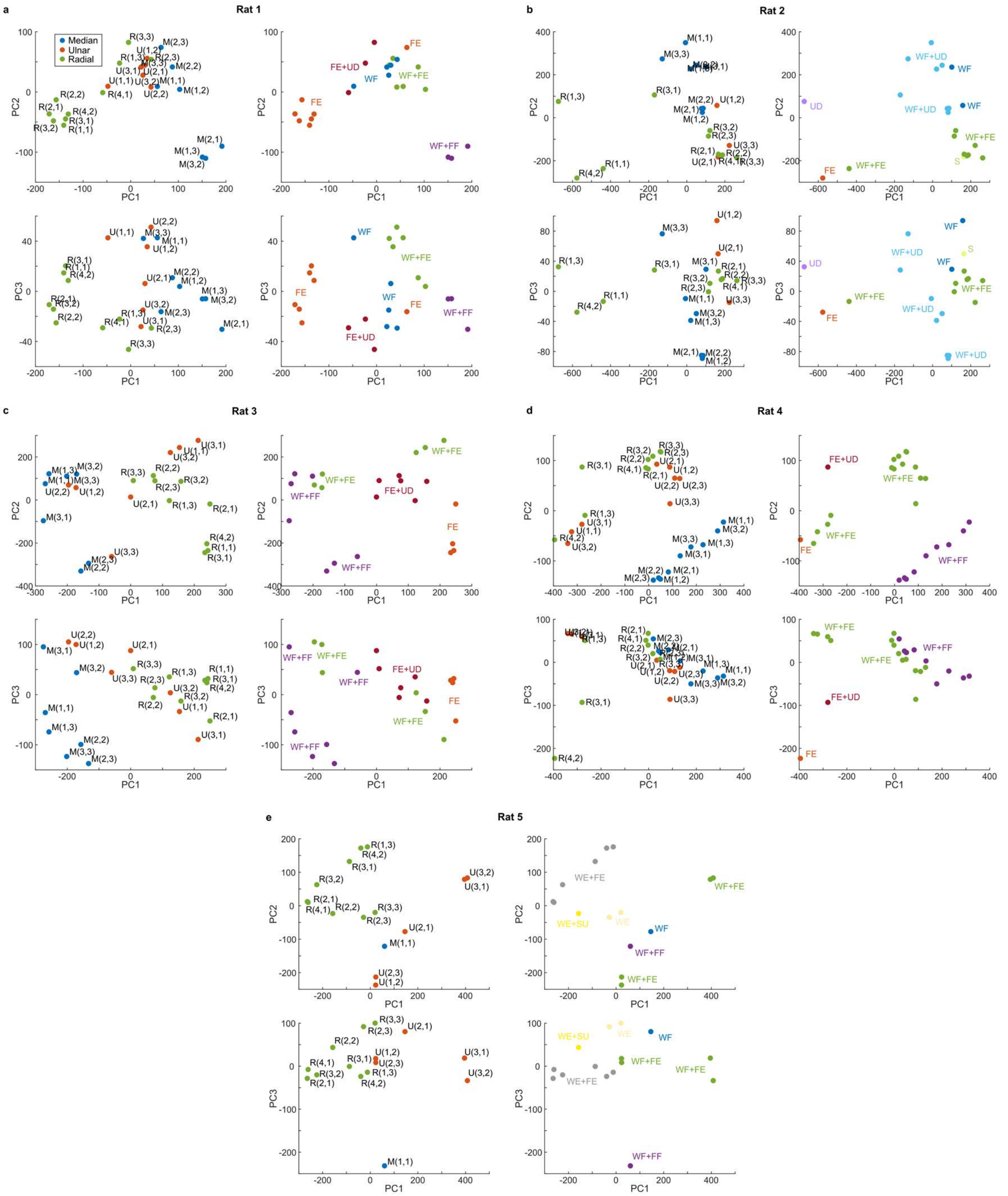


**Supplementary Fig. 7 | Analysis of kinematics of movements induced by nerve stimulation through ultraconformable cuffs. a-e**, Principal component analysis of kinematics for Rat 1 (**a**), Rat 2 (**b**), Rat 3 (**c**), Rat 4 (**d**) and Rat 5 (**e**). Top panels: plots of principal components 1 against 2. Bottom panels: plots of principal components 1 against 3. Left panels: movements are tagged based on the microelectrode that produced them. Right panels: movements are tagged by the identified type of movement produced. WF: wrist flexion, WE: wrist extension, FE: finger extension, UD: ulnar deviation, S: supination, WF+FF: wrist flexion and finger flexion, WF+FE: wrist flexion and finger extension, WF+UD: wrist flexion and ulnar deviation, WE+FE: wrist extension and finger extension, FE+UD: finger extension and ulnar deviation. Bottom panels in a) correspond to those of Fig. 5e-f.

**Supplementary Video 1| Examples of other types of paw movements produced through stimulation with two microelectrodes simultaneously.** First (Finger extension + Wrist flexion) and second (Wrist extension + Wrist flexion) clips correspond to a one rat, while the third (Finger extension + Finger flexion) corresponds to a different rat. Microelectrodes delivering stimulation are depicted in red in implant diagram.
